# Tracking attention in a visual active paradigm for the diagnosis of disorders of consciousness

**DOI:** 10.1101/2019.12.11.872515

**Authors:** Damien Lesenfants, Camille Chatelle, Steven Laureys, Quentin Noirhomme

## Abstract

**Background:** Clinical assessment of patients with disorders of consciousness (DOC) relies on the clinician’s ability to detect a behavioral response to an instruction (e.g., “squeeze my hand”). However, recent studies have shown that some of these patients can produce volitional brain responses to command while no behavioral response is present. This highlights the importance of developing motor-independent diagnostic tool for this population, complementing standardized behavioral evaluation. We here evaluate the ability of a novel gaze-independent attention-based EEG paradigm to detect volitional attentional processes in patients with disorders of consciousness.

**Methods:** Thirty patients with DOC were included in the study: 12 with an unresponsive wakefulness syndrome, 16 in a minimally conscious state (MCS), two who emerged from a MCS. Patients were randomly instructed to either concentrate on a task or rest while brain activity was recorded using EEG during a gaze-independent paradigm.

**Results:** One of two EMCS, one of 16 MCS and one of 12 UWS patients showed a response to command using the attention task. Interestingly, this method could detect a brain-based response to command in one MCS patient who did not present a behavioral response to command at the bedside the day of the assessment.

**Conclusion:** This study show that task-related variation of attention during an active task could help to objectively detect response to command in patients with DOC.

## Introduction

Holy Grail in the diagnosis of patients with disorders of consciousness (DOC) is the detection of a volitional and goal-oriented response to command. Indeed, “command following”, the ability to execute a simple command, is a clinical landmark distinguishing patients with a unresponsive wakefulness syndrome (UWS; characterized by the recovery of eye opening without awareness; Laureys et al. 2010) from patients in a minimally conscious state (MCS; showing inconsistent but reproducible behaviors and unable to communicate; Giacino et al. 2002). Current clinical assessment of the patient’s conscious state relies on an examiner’s detection of behavioral responses to command based on visual (e.g., “follow my finger movement with your eyes”) and tactile (e.g., “squeeze my hand”) feedback using behavioral scales, like the Coma Recovery Scale-Revised (CRS-R; Giacino et al. 2004).

In 2006, researchers from Universities of Liege and Cambridge showed a consistent response to command, independent of any motor pathway, with functional magnetic resonance imaging in a patient diagnosed as being in a vegetative/unresponsive wakefulness syndrome (Owen et al. 2006). This patient was instructed to imagine “playing tennis” and “walking through her house” during the acquisition of functional magnetic resonance imaging-based brain response to task. In a follow up study including 54 patients (Monti et al. 2010), five were able to willfully modulate their brain activity. One of them was even able to answer simple questions using one task for “yes” and the other for “no”. This study suggested that the absence of responsiveness does not necessarily correspond to absence of awareness (Sanders et al. 2012), illustrating the need to develop motor-independent diagnostic and communication tools for this population suffering from limited neuromuscular abilities (Laureys & Schiff 2012; Stender et al. 2014), allowing the objective detection of covert volitional response to command at bedside and complementing standardized behavioral evaluation. Moreover, behavioral assessment requires trained experienced caregivers and can suffer from the examiner’s subjectivity (Løvstad et al. 2010). Improving diagnoses and unlocking communication ability in these patients could lead to improved rehabilitation strategies, quality of life, and prognosis (e.g., an early recovery of a response to command is associated with a more favorable outcome, Whyte et al. 2009).

Building upon this work, recent research has investigated patients’ abilities to generate EEG responses to different modalities, removing the limited availability in clinics and patient’s motion sensitivity of the functional magnetic resonance imaging technology. Visual, motor, somatosensory, auditory or hybrid (i.e., integrating multiple stimulation paths) modalities have been investigated in DOC patients (for a review, see Lesenfants et al. 2016). However, despite the increasing interest of the community in working with DOC patients, high false-negative (i.e., the percentage of patients showing a response to command behaviorally but not with the system; currently ranging from 25% to 100%, see Chatelle et al. 2015) and false-positive (i.e., the percentage of unconscious patients wrongly showing a response to command with the system) rates illustrate the difficulty in translating this technology from healthy individuals to severe brain-injured patients.

Recently, we proposed a novel gaze-independent attention-based EEG paradigm allowing the detection of volitional attentional processes (“concentrate versus rest”) with 93% and 91% accuracy in healthy volunteers and patients with a locked-in syndrome, respectively (performance at chance level in unconscious patients) (Lesenfants et al. 2018). Here, we used this approach in a broader group of patients with DOC in order to assess its potential as a diagnostic tool for this population.

## Methods

A convenience sample of 30 DOC patients were included: 12 patients with an unresponsive wakefulness syndrome (8 men; age, mean ± SD, 43 ± 15 years; mean time post injury, mean ± SD, 32 ± 26 months; 10 patients were already part of a previously published study Lesenfants et al. 2018), 16 with a minimally conscious state (8 men; age, 42 ± 13 years; mean time post injury, 61 ± 85 months; 11 with a behavioral response to command, MCS+) and two who emerged from the minimally conscious state (EMCS; age, 24 ± 14 years; mean time post injury, 18 ± 11 months). The study was approved by the Ethics Committee of the University Hospital of Liège, and all participants or their legal representative provided both informed and written consent. Patients were assessed by a trained examiner using the CRS-R on the day of the EEG recording and several times during the week to increase diagnostic accuracy (at least 5 times; Giacino et al. 2004). The best score obtained during the week was used as the final diagnosis. Occipital activity was recorded using 12 Ag/AgCl ring electrodes at location P3, P1, P2, P4, PO7, PO3, POz, PO4, PO8, O1, Oz and O2, referenced to Pz, based on the international 10-20 electrode system. Ground electrode was placed behind the right mastoid. All impedances were kept below 5 kΩ. The amplifier used was a BrainVision V-Amp with a band pass filter between 0.01 and 100 Hz and a sampling frequency of 250 Hz. A visual stimulation pattern composed of flickering yellow (10 Hz) and red (14 Hz) interlaced light emitting diode-squares was placed at 30 cm of the patient’s head (Lesenfants et al. 2014). The session was composed of six blocks, each lasting 5 minutes. During a block, we requested the patient to actively focus on one of two colors (active trials, 5 red vs 5 yellow randomly presented 7s-trials) or to rest (passive trials, 10 7s-trials). Inter-trials interval were used to deliver the auditory instructions via the headphones. Seven seconds band-pass (Butterworth 4th order; 5-60 Hz) and band-stop (Notch IIR; 50Hz) filtered EEG data were used as a unique window for each active and passive trials. Steady-state visually evoked responses (i.e., the power at the first harmonic of each stimulation frequency, Lesenfants et al. 2014) were extracted on each active trials using a Multitapers Spectral Analysis (7 tapers; Thomson 1982). Normalized entropy measures (Viertio-Oja et al. 2004) were extracted from active and passive trials using power spectral estimation based on Multitapers Spectral Analysis (7 tapers; frequency band used is 0.5-32 Hz with a 0.5Hz step). Visual (active red versus active yellow) and attentional (active versus passive) classification performances were then computed with a linear discriminant analysis (LDA), and assessed with a 10×10-fold cross-validation. A permutation test (Nichols & Holmes 2002; Noirhomme et al. 2017) evaluated the chance-level for each participant (1000 repetitions, LDA classification, 10×10-fold cross validation, p<0.01). The significance of change between conditions was assessed with a non-parametric Wilcoxon signed-rank test (2-tailed, p < 0.01), with Bonferroni correction for multiple comparisons. All analyses were done with custom made code using Matlab. The BCI2000 software package (Schalk et al. 2004) and Fieldtrip Toolbox (Oostenveld et al. 2011) were used for data acquisition and presentation of the auditory instructions.

## Result

One of two EMCS (accuracy: 69 ± 2%), one of 16 MCS (64 ± 2%) and one of 12 UWS (67 ± 2%) patients showed a response to command (permutation test, p < 0.01) using the attention task. None of the patients with DOC showed a response to command (i.e., a classification accuracy above chance level) using the visual task.

A decrease of the EEG power at the red stimulation frequency (14Hz) between the passive trials and the active trials could be observed for the MCS+ group when requesting to focus on the red flashing squares (WSRT, p < 0.01; see Figure 1, upper left panel). When requesting to focus on the yellow flashing squares, a decrease of the EEG power at the yellow stimulation frequency (10Hz) between the passive trials and the active trials could be observed for the EMCS (WSRT, p < 0.01; see Figure 1, lower left panel).

**Fig. 1.**
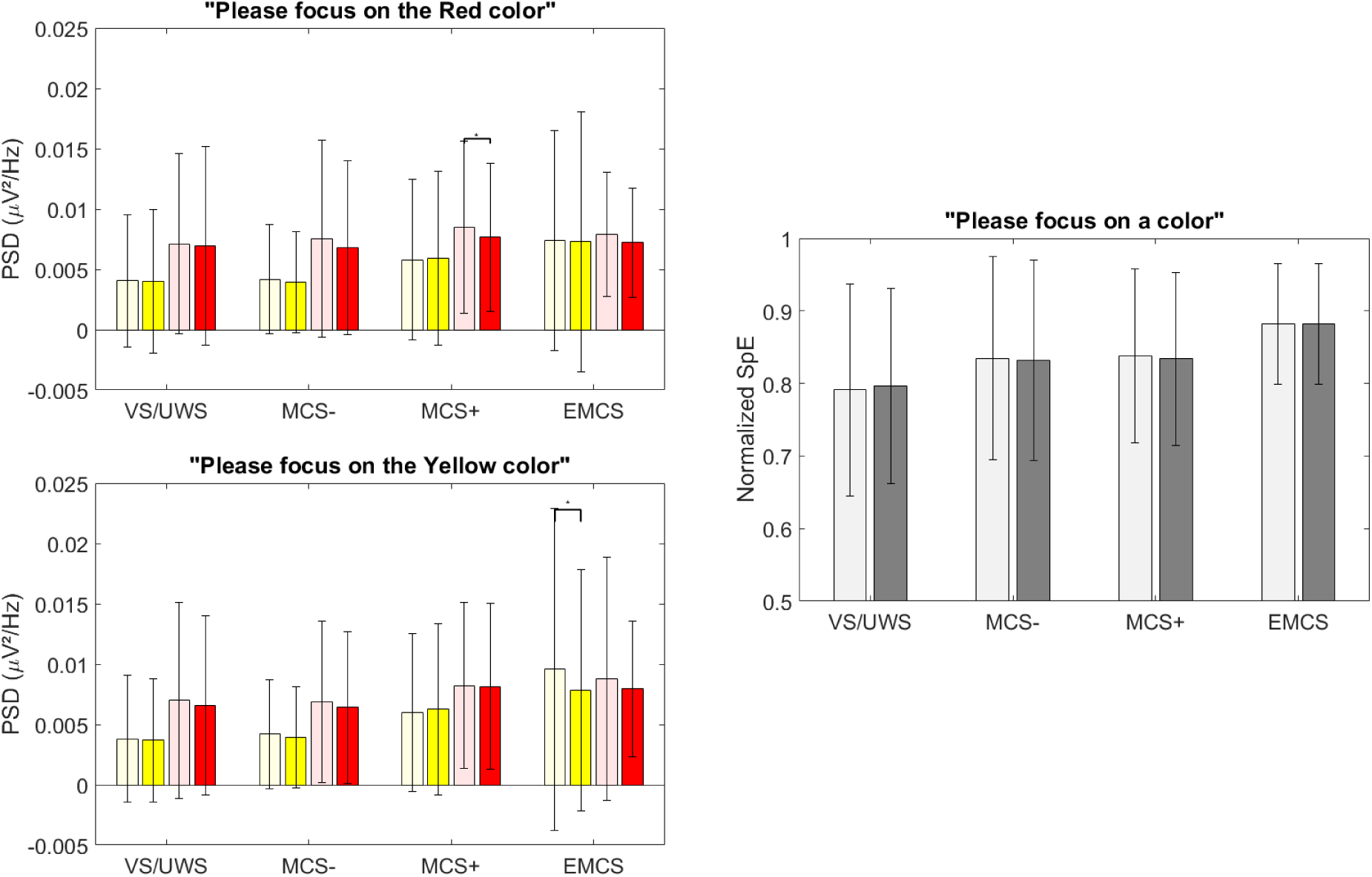
Left: Amplitude of the 10Hz (in yellow) and 14Hz (in red) SSVEP response (multitapers, 7 tapers) over the 7s-active red (upper panel) and yellow (lower panel) trials as well as the preceeding 7s-resting periods (transparent) for the different patient’s group. Right: Averaged spectral entropy during the 7s-active (dark gray) and 7s-passive (light gray) periods.

The averaged spectral entropy, computed during either the passive or the active trials, increased from VS to MCS (and EMCS; see Figure 1, right panel).

At single-subject level, the spectral entropy increased from the passive periods to the active periods and the related instruction (“please focus on a color”) for patients VS/UWS6 and EMCS2 (WSRT, p < 0.01; see Figure 2).

**Fig. 2.**
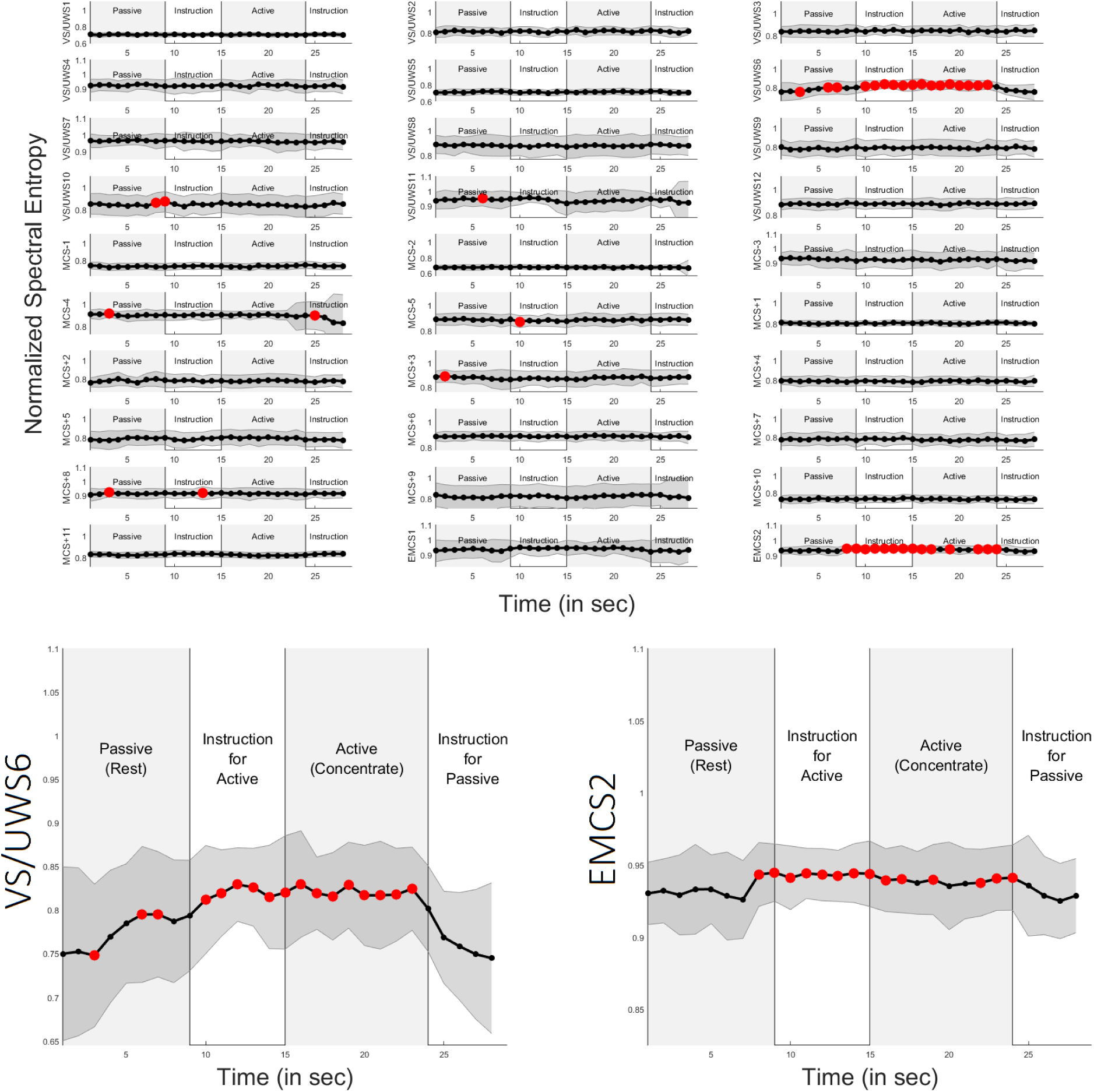
Individual spectral entropy computed over consecutive 1s-periods during the 30s active vs passive trials and averaged over trials. Red dots represents 1s-periods with significantly (WSRT; p < 0.01) higher spectral entropy as compared to the passive 7s-periods (rest). Note the increase of the spectral entropy during the active periods and the related instruction (“please focus on a color”) for patients VS/UWS6 and EMCS2.

## Discussion

The proposed method allowed detecting a brain-based response to command in three patients. One of them, finally diagnosed as MCS+ (i.e., showing minimal signs of consciousness and able to produce a response to command), did not present a behavioral response to command at the bedside the day of the assessment. This shows the importance in complementing behavioral evaluation with objective diagnostic tool. It is important to stress that, while we hope that this technology could a day replace the behavioral evaluation, results still need to be confirmed in a multi-site study allowing for an extended cohort of patients. To this day, the gold standard is the behavioral scales and novel tools should only complement their results by providing extra information. In the specific scenario in which a patient presents a brain-based response to command but no sign of consciousness at the bedside, a follow-up should be consider to evaluate if the neuroimaging tool provides a false-positive or corrects for a behavioral false-negative (Monti et al. 2010; Stender et al. 2014; King et al. 2013; Schnakers et al. 2009; Chennu et al. 2013).

At group-level, we observed an increase of the averaged spectral entropy with patient’s level of consciousness similarly to Gosseries et al. (2011). This could be partially explained by a slowing down of the EEG with patient’s condition, more marked in UWS than in MCS patients (Lehembre et al. 2012). Interestingly, the amplitude of the EEG response at the target stimulation frequency, i.e. the EEG amplitude at the frequency of the red (resp. yellow) stimuli when requesting the patient to focus on the red (resp. yellow) pattern, decreased between the passive and the active periods in MCS+ and EMCS groups. This could be due to the covert stimulation involving an inhibitory mechanisms in the brain act to suppress responses to distracting stimuli (Kastner & Ungerleider 2000).

Two patients (VS/UWS6 and EMCS2) showed a significant increase of the spectral entropy from the passive to the active periods (see Figure 2), suggesting a volitional modulation of the attentional level with the task. These patients also present a brain-based response to command using machine learning based brain pattern classification.

Despite its gaze-independency, only one out of the five patients (see EMCS2) showing a behavioral response to command at the bedside the day of the assessment showed a response to command with our method. This illustrates the complexity in detecting a volitional response to command in patients with DOC using an EEG-based diagnostic tool (based on the final clinical diagnosis: false negative rate: 85%, false positive rate: 8%). This could be due to the complexity and the mental demand of this paradigm. We could also hypothesize that the non-stationarity in the EEG signals and patients’ fluctuation of level of consciousness limit the application of automatic EEG-pattern classification in this context. Future research should consider a fully adaptive classification process, adapting parameters at single-trial level, and tracking attention during a simpler and shorter task.

## Acknowledgements

We are grateful to the patients and their families for participating in the study. The study was supported by the University and University Hospital of Liège, the French Speaking Community Concerted Research Action (ARC 12-17/01), the Belgian National Funds for Scientific Research (FRS-FNRS), Human Brain Project (EU-H2020-fetflagship-hbp-sga1-ga720270), Luminous project (EU-H2020-fetopen-ga686764), the James McDonnell Foundation, Mind Science Foundation, IAP research network P7/06 of the Belgian Government (Belgian Science Policy), the European Commission, the BIAL Foundation, the Public Utility Foundation Université Européenne du Travail, Fondazione Europea di Ricerca Biomedica, Belgian National Plan Cancer (139), ECH2020 project ComaWare. CC is a Marie Sklodowska-Curie fellow (H2020-MSCA-IF-2016-ADOC-752686).

---

**Table 1.** Demographic, clinical and task-related data of the patients’ sample. Onset column indicate the interval since insult (in months). Behavioral Assessment columns indicate the CRS-R subscores at the day of the EEG assessment for respectively Auditory, Visual, Motor, Verbal, Communication and Arousal functions, and related diagnosis. Visual column illustrate the “active red versus active yellow” classification accuracy (in percent) obtained with the multi-trial steady-state visually evoked potential analysis. Attention column illustrates the “active versus passive” classification accuracy (in percent) obtained with the multi-trial spectral entropy analysis. Multi-trial accuracies above chance-level (permutation test, p<0.01, 1000 permutations) are highlighted with an asterisk (*).

|  | Age | Etiology | Onset | Behavioral Assessment |  | Visual | Attention |
| --- | --- | --- | --- | --- | --- | --- | --- |
|  |  |  |  | Score | Diagnosis |  |  |
| UWS1 | 60-65 | Trauma | 7 | 1-1-2-1-0-2 | UWS | 57±5 | 54±2 |
| UWS2 | 20-25 | Trauma | 10 | 1-1-2-1-0-1 | UWS | 65±6 | 56±3 |
| UWS3 | 50-55 | Trauma/anoxia | 10 | 1-1-2-1-0-1 | UWS | 49±5 | 51±6 |
| UWS4 | 45-50 | Subarachnoid | 12 | 1-0-1-2-0-1 | UWS | 49±9 | 50±3 |
| UWS5 | 50-55 | Subarachnoid | 14 | 1-0-1-1-0-1 | UWS | 44±4 | 46±4 |
| UWS6 | 50-55 | Anoxia | 17 | 1-1-1-1-0-1 | UWS | 41±5 | **67±2 (\*)** |
| UWS7 | 60-65 | Cardiac arrest | 26 | 1-0-2-0-0-1 | UWS | 53±5 | 48±3 |
| UWS8 | 30-35 | Anoxia | 36 | 1-0-2-1-0-1 | UWS | 67±7 | 49±4 |
| UWS9 | 20-25 | Anoxia | 43 | 1-0-1-1-0-2 | UWS | 47±6 | 47±4 |
| UWS10 | 35-40 | Cardiac arrest | 57 | 1-0-1-1-0-2 | UWS | 54±3 | 47±1 |
| UWS11 | 30-35 | Trauma | 65 | 1-0-1-1-0-1 | UWS | 56±5 | 48±2 |
| UWS12 | 25-30 | Trauma | 86 | 1-0-1-2-0-1 | UWS | 56±2 | 48±4 |
| MCS-1 | 40-45 | Trauma | 8 | 1-0-2-1-0-1 | UWS | 54±5 | 38±4 |
| MCS-2 | 25-30 | Trauma | 18 | 1-2-2-1-0-1 | MCS- | 46±6 | 47±3 |
| MCS-3 | 50-55 | Cardiac arrest | 49 | 1-1-2-2-0-1 | MCS- | 61±6 | 54±3 |
| MCS-4 | 30-35 | Trauma | 105 | 1-1-2-2-0-1 | UWS | 49±4 | 47±3 |
| MCS-5 | 40-45 | Trauma | 359 | 0-3-2-2-0-2 | MCS- | 32±5 | 44±4 |
| MCS+1 | 60-65 | Trauma | 4 | 1-1-2-1-0-2 | MCS- | 49±6 | 46±6 |
| MCS+2 | 45-50 | Cardiac arrest | 7 | 1-0-1-1-0-1 | UWS | 47±4 | 48±2 |
| MCS+3 | 45-50 | Trauma | 11 | 1-1-2-2-0-1 | MCS- | 45±4 | 45±3 |
| MCS+4 | 65-70 | Anoxia | 11 | 0-3-2-2-0-2 | MCS- | 39±3 | 40±4 |
| MCS+5 | 30-35 | Subarachnoid | 19 | 3-4-3-2-0-2 | MCS+ | 55±5 | 44±2 |
| MCS+6 | 40-45 | Trauma | 21 | 0-4-2-2-0-1 | MCS- | 47±4 | 46±2 |
| MCS+7 | 20-25 | Trauma | 36 | 0-3-2-1-0-1 | MCS- | 38±8 | 53±4 |
| MCS+8 | 30-35 | Trauma | 40 | 3-1-2-1-0-1 | MCS+ | 58±5 | 42±3 |
| MCS+9 | 25-30 | Trauma | 61 | 3-4-5-1-0-2 | MCS+ | 47±8 | 57±3 |
| MCS+10 | 30-35 | Trauma | 90 | 0-0-2-1-0-2 | UWS | 41±5 | 60±2 |
| MCS+11 | 55-60 | Trauma | 136 | 2-3-1-1-0-2 | MCS- | 44±5 | **64±2 (\*)** |
| EMCS1 | 10-15 | Subarachnoid | 6 | 3-5-6-1-1-1 | EMCS | 56±5 | 46±3 |
| EMCS2 | 30-35 | Trauma | 29 | 3-5-2-1-0-1 | MCS+ | 49±4 | **69±2 (\*)** |

